# Divergent outcomes of direct conspecific pathogen strain interaction and plant co-infection suggest consequences for disease dynamics

**DOI:** 10.1101/2022.11.01.514679

**Authors:** Hadjer Bellah, Nicolas F. Seiler, Daniel Croll

**Affiliations:** Laboratory of Evolutionary Genetics, Institute of Biology, University of Neuchâtel, CH-2000, Neuchâtel, Switzerland

**Keywords:** fungi, competition, virulence, plant pathogens, co-infection, Zymoseptoria tritici

## Abstract

Plant diseases are often caused by co-infections of multiple pathogens with the potential to aggravate disease severity. In genetically diverse pathogen species, co-infections can also be caused by multiple strains of the same species. However, the outcome of such mixed infections by different conspecific genotypes is poorly understood. The interaction among pathogen strains with complex lifestyles outside and inside of the host are likely shaped by diverse traits including metabolic capacity and the ability to overcome host immune responses. To disentangle competitive outcomes among pathogen strains, we investigated the fungal wheat pathogen *Zymoseptoria tritici*. The pathogen infects wheat leaves in complex strain assemblies and highly diverse populations persist between growing seasons. We investigated a set of 14 genetically different strains collected from the same field to assess both competitive outcomes under culture conditions and on the host. Growth kinetics of co-cultured strains significantly deviated from single strain expectations indicating competitive exclusion depending on the strain genotype. We found similarly complex outcomes of lesion development on plant leaves following co-infections by the same pairs of strains. While some pairings suppressed overall damage to the host, other combinations exceeded expectations of lesion development based on single strain outcomes. Strain competition outcomes in absence of the host were poor predictors of outcomes on the host suggesting that the interaction with the plant immune system adds significant complexity. Intraspecific co-infection dynamics likely make important contributions to disease severity and need to be integrated for a more complete understanding of host-pathogen dynamics in the environment.

**Importance:** Plants are often attacked by a multitude of pathogens simultaneously. Different pathogen species can either facilitate or constrain the colonization by other pathogen species. Hence, natural infections are often the outcome of complex interactions between pathogens. To what extent the simultaneous colonization of genetically different strains of the same pathogen species matters for disease outcomes remains largely unclear though. We assessed the outcome of interactions between strains of the fungal wheat pathogen Zymoseptoria tritici. In absence of the host, strains cultured in pairs were growing differently compared to strains cultured alone. When infecting wheat leaves either with single or pairs of strains, we found also highly variable outcomes. Importantly, interactions between strains outside of the host were only poorly explaining how strains would interact when on the host. This suggests that pathogen strains engage in complex interactions shaped by their environment. Understanding the nature of such interactions within pathogen species will improve our ability to manage crop plant infections in the wild.

## Introduction

Plants are exposed to a variety of microorganisms in their natural environment (1–3). Colonizing microorganisms often occur as complex communities with complex interkingdom (mainly bacteria and fungi) interactions on single plant hosts (4–7). Colonization by multiple pathogenic organisms are knowns as co-infection and are common in nature (5). Co-infections are of particular interest since the interaction between pathogens can significantly alter infection severity (8). Because coinfections can also modulate the selection pressure on pathogens, the trajectory of virulence evolution can be modulated by co-infections (9, 10). Co-infecting pathogens can cause varying levels of damage to the host depending on the outcome of microbial interactions and host immune responses (11). For example, previous studies showed that when plants are co-infected by pathogens of different species, one species tends to suppress the growth and development of the other competing species (12). However, at least some co-infections seem to have no detectable impact on disease symptoms compared to single infections.

The outcome of co-infections can be affected by the order of co-infection. Sequential co-infection can decrease the virulence of the second strain involved in the co-infection because of the priming of plant defenses (13), however such effects can be genotype dependent (14). The host immune defenses activated by the first pathogen might not be mounted rapidly enough to prevent damage, however the second infecting pathogen may be limited by primed immune defenses (15). Consistent with this phenomenon, simultaneous infections can be more harmful to hosts than sequential infections (16). Co-infections increasing disease severity through pathogen cooperation were observed, for example, for microbial cooperation increasing disease caused by the ascomycete *Didymella bryoniae* by helping to translocate four bacterial species that co-infect Styrian oil pumpkin (17). Competition among individuals of the same species can also occur (18) with conflicts over access to nutrients in well-defined niches such as plant leaves (19). Interactions between co-infecting pathogens are context-dependent with many factors influencing the outcome of infections and are, hence, difficult to predict.

For plants, intraspecific co-infections are playing a key role in disease progression with fungal-plant pathosystems receiving most attention (5, 20, 21). Plant pathogenic fungi display a wide range of specialization levels from generalists able to infect hundreds of host plants to specialists, which are infectious only on single host species (22–24). Within pathogen species, strains can show a high degree of variation in virulence (20, 21). The virulence of co-infecting *Podosphaera plantaginis* strains can be increased on the *Plantago lanceolata* host (5). In other systems such as *Zymoseptoria tritici* infecting wheat, virulence appears unaffected by co-infections (20, 21), however the trajectory of an infection depends on the host genotype and genotypes of competing strains (20, 25). Besides, the competitive advantage of individual strain genotypes for transmission is not associated with virulence among the strains (25). The growth of mixed genotypes outside of the host is only poorly associated with the success of the same genotype mix on the plant though (25). The complex links between competitive abilities of the pathogen and consequences for the plant host raise important questions how co-infections impact evolutionary trajectories of host-pathogen systems.

*Z. tritici* is a major fungal pathogen of wheat causing the economically important foliar disease Septoria tritici blotch (STB) (26). *Z. tritici* exhibits a symptomless phase after coming into contact with wheat leaves lasting 8–11 days post infection (dpi), during which the fungus does not induce visible host defense responses (27, 28). In later stages, the fungus invades the intercellular space of leaves producing lesions and asexual fruiting bodies called pycnidia. Under co-infection, *Z. tritici* strains colonize wheat leaves in a mosaic of genotypes (29). Field populations exhibit high genetic and phenotypic diversity worldwide due to frequent sexual reproduction, large populations, and significant gene flow among populations (30–32). Co-infection of wheat leaves by multiple strains of *Z. tritici* are frequent in the field (29).

The aim of this study was to investigate the outcome of pathogen strain co-existence both outside of the host and on the host. For this, we analyzed 15 different strains sampled from the same wheat field and set up 91 and 105 different pairs to assess competitive outcomes among strains both inside and outside of the host respectively and the impact of co-infection on disease severity.

## Results

### In-vitro growth rate assessment and divergent outcomes of co-cultures

We analyzed the growth of 14 *Z. tritici* strains individually and in 91 pairwise combinations in liquid growth medium (See Supplementary Figure S1; Supplementary Tables S1-2). In general, we observed stationary growth after 11 days of incubation at 18 °C (Fig. 1A). We assessed different culture growth metrics including the maximum specific growth rates (*μmax*) and doubling times. Maximum specific growth rates (*μmax*) and doubling times for different strains were highly correlated (Pearson correlation, *r* = −0.96, *p-value* < 0.0001; Fig. 1B). Single strains displayed variable maximum specific growth rates (*μmax*) (Fig. 1C). We investigated the overall growth in co-cultures compared to the growth of individual strains used to establish the co-culture. Analyzing the 91 pairs established as co-cultures, we found that 37 co-cultures showed a significant difference in growth compared to strains grown independently (*p-value* < 0.05; Fig. 1D). Each of the 14 strains used for the pairings was showing a growth significantly different in at least one pairing (*p-value* < 0.05; Fig. 1D). The strain 3K8 showed significantly higher maximum specific growth rates (*μmax*) in 9 out of the 13 pairings (*p-value* < 0.05; Fig. 1D). Strain 1S10 was growing significantly faster in 6 out of 13 pairings (*p-value* < 0.05; Fig. 1D). In contrast, the strains 1G30, 1P35, 3R39, 3W21 and 3W3 were significantly slower growing compared to all their respective pairings (*p-value* < 0.05; Fig. 1D). We found that 7 out of 91 co-cultures were growing not significantly different to one of the two strains to establish the mixture while the other strain grew significantly less alone (Fig. 2A; pattern a; for examples See Supplementary Figure S2). Furthermore, 17 out of 91 pairings showed the opposite pattern with one of the strains in the pairing growing significantly more alone compared to the mixture (Fig. 2A; pattern b; for examples See Supplementary Figure S2). Additional patterns included either one or both strains growing alone significantly less than the co-culture (Fig. 2A; pattern c and f; for examples See Supplementary Figure S2). We also observed 7 co-cultures growing with no significant differences to both single strain cultures (Fig. 2A; pattern d; for examples See Supplementary Figure S2). Besides, we observed 7 co-cultures growing at an intermediate level with significant differences to both single strain cultures (Fig. 2A; pattern g; for examples See Supplementary Figure S2). Finally, we observed no significant difference in growth among either single strains or the co-culture for 38 out of 91 interactions (Fig. 2A; pattern e; for examples See Supplementary Figure S2).

**Figure 1:**
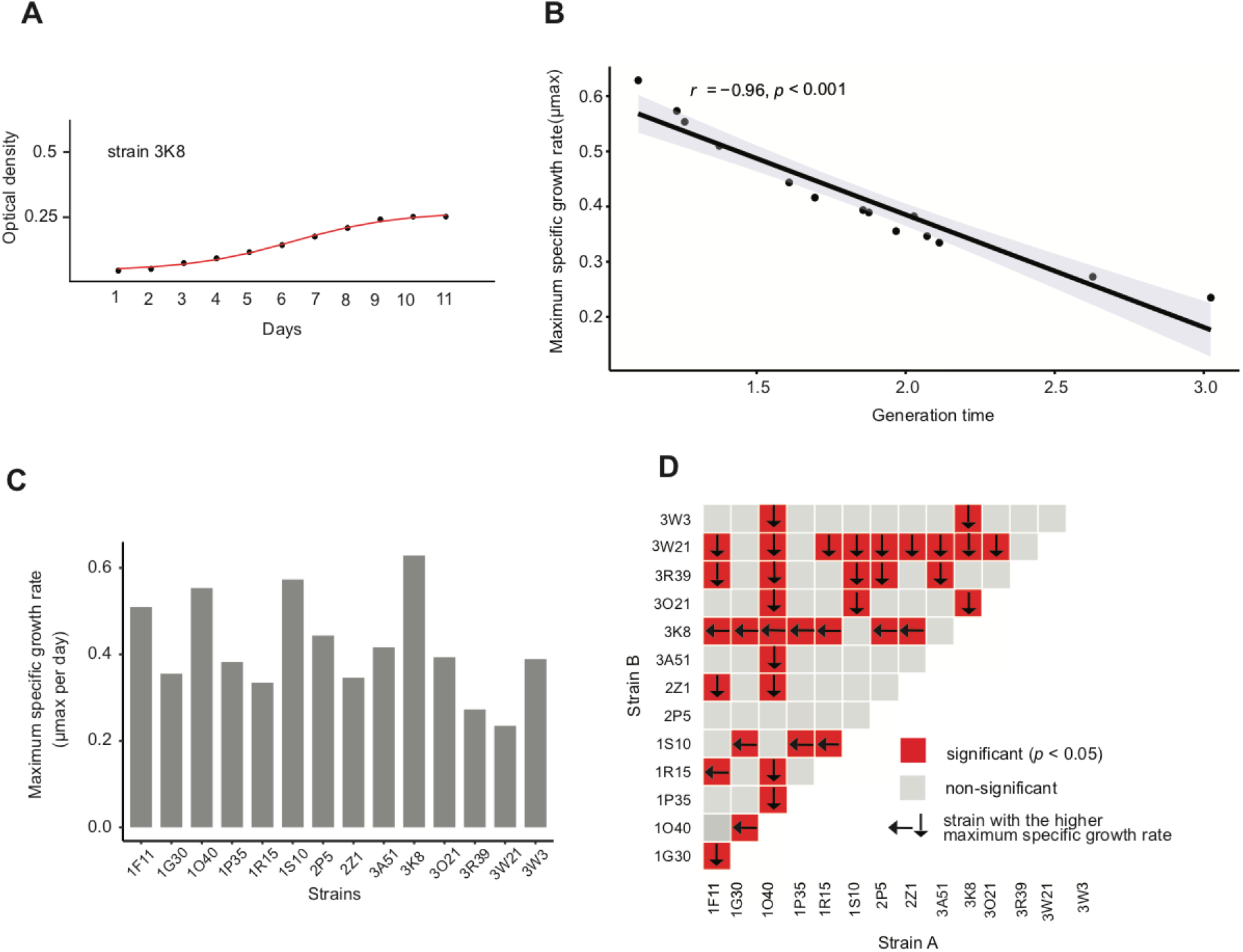
Culture condition growth of *Zymoseptoria tritici* strains. A) Growth curve of optical density during the lag, exponential and stationary phase. A growth plateau was typically reached after ~11 days of incubation in the dark at 18 °C. B) Correlation of growth metrics including the maximum specific growth rate (*μmax*) and doubling time across strains (Pearson correlation, r = −0.96, *p-value* < 0.0001). C) Maximum specific growth rate (μmax) variation among different strains. D) The growth of strains involved in different pairwise interactions assays. Red indicates significant differences between growth rates of individual cultures. The arrow points towards the strain with the faster growth rate.

**Figure 2:**
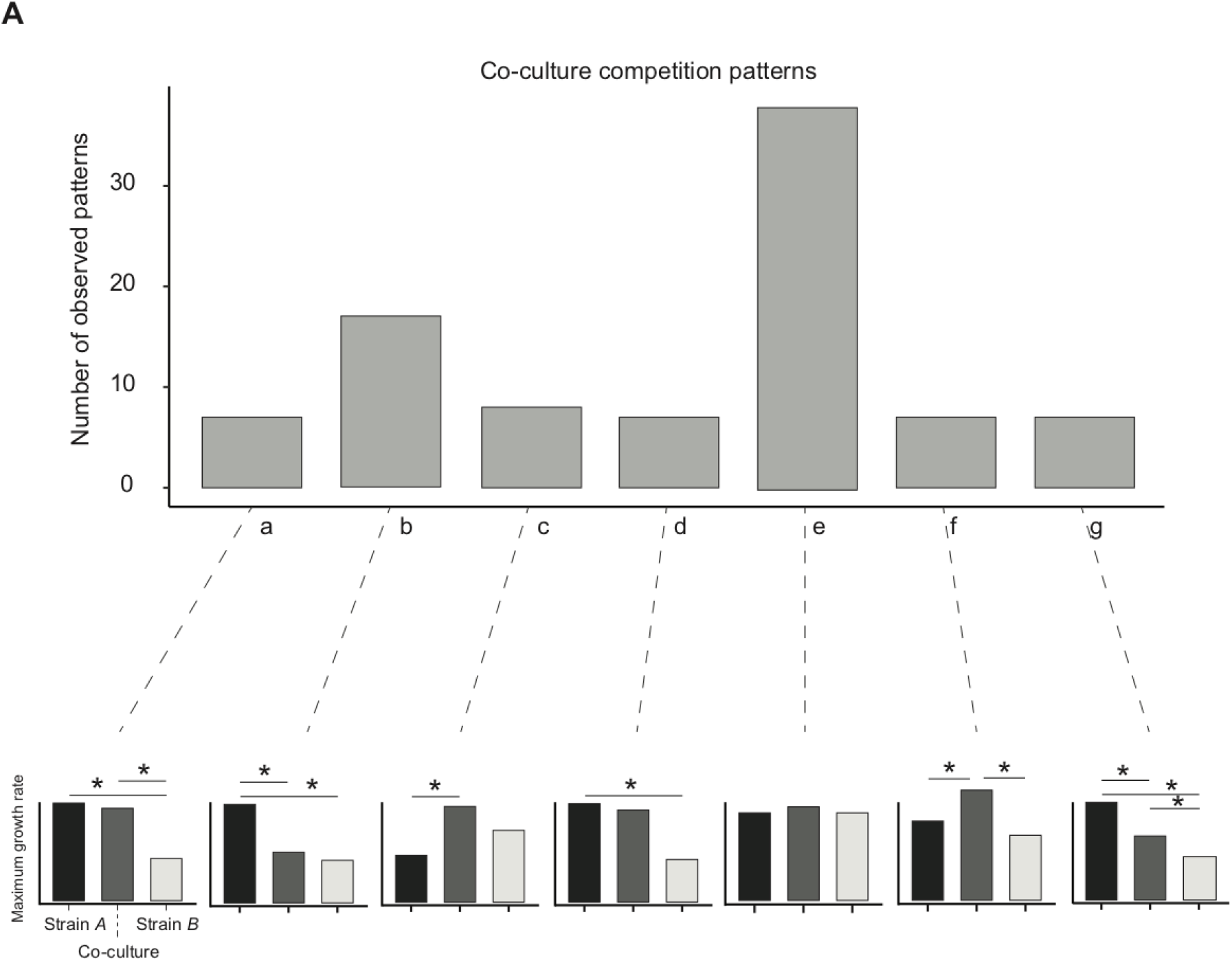
Single and co-culture outcomes observed for the *in vitro* experiments. All interactions were grouped into seven types of pairwise interaction patterns (letters a-g).

### Variation in outcomes among single and co-infections of wheat

We analyzed infection symptoms on wheat leaves produced both by single strain infections (*n* = 15) and 105 pairs created from the same set of strains. Disease symptoms were assessed as the percent leaf area covered by lesions (PLACL). The measurements were made using scans of entire wheat leaf and contrast-based image analysis (Fig. 3A; See Supplementary Table S4). Single strain infections produced variable disease symptoms from PLACL ranging from 0-70% (Fig. 3B). For the 105 different pairs, ten pairs included strains with a significant difference in PLACL when infected alone (*p-value* < 0.05; Fig. 3C). The ten pairs included pairings of a total of ten different strains. We analyzed the outcomes of the co-infection compared to the single infections similarly to the culture conditions. We observed two and four out of 105 pairs with one strain being either significantly lower or higher, respectively, compared to the other strain and the co-infection (Fig. 4A-B; patterns *a* and *b*). We observed higher numbers of interactions where either one strain alone was producing significantly less symptoms compared to the co-infection or compared to the other strain (Fig. 4A-B; patterns *c* and *d*). Over two thirds (72 out of 105) of all interactions tests showed no significant differences between either strain alone or the co-infection (Fig. 4A-B; pattern *e*). Finally, the last relevant virulence pattern consisted of two strains, which were not statistically different, but the co-infection was significantly different from the two strains infecting individually the plant (*p-value < 0.0*5; Fig. 4A-B; patterns *f*). We found that 7 out of 105 combinations followed this pattern (Fig. 4A-B; pattern *f*). Interestingly, in all tested combinations following this latter pattern, the virulence of co-infection was always lower than observed during single infections.

**Figure 3:**
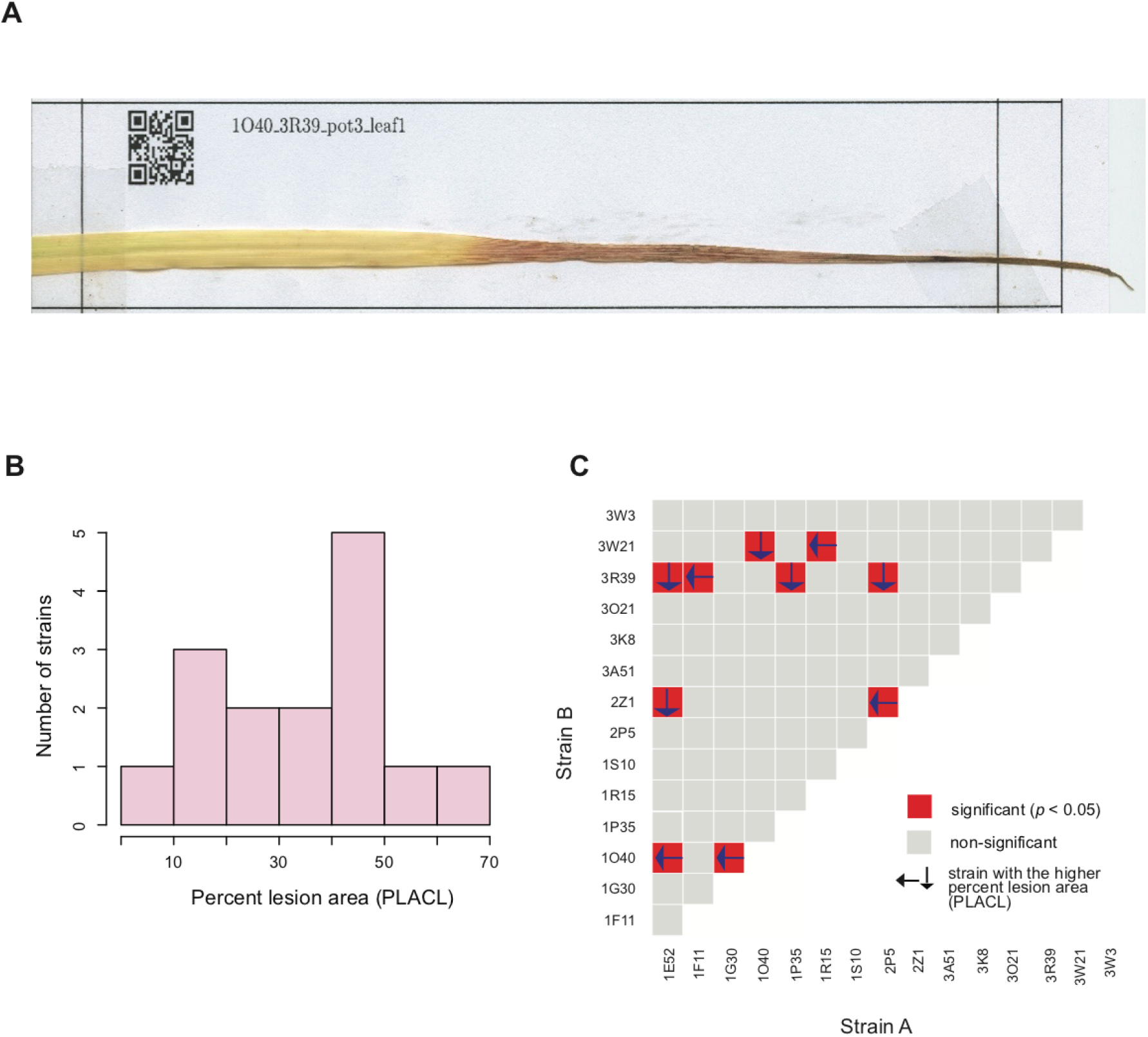
Infection outcomes of single and co-infections of wheat leaves. A) Scan of a mounted wheat leaf at the end of the infection period. The extent of the lesion area over the entire leaf was assessed. B) Distribution of lesion area produced by each strain in a single infection experiment. C) Assessment of co-infection outcomes of all pairs. For significant differences among co-culture and single infections, the strain producing the larger lesion area when alone is highlighted by an arrow.

**Figure 4:**
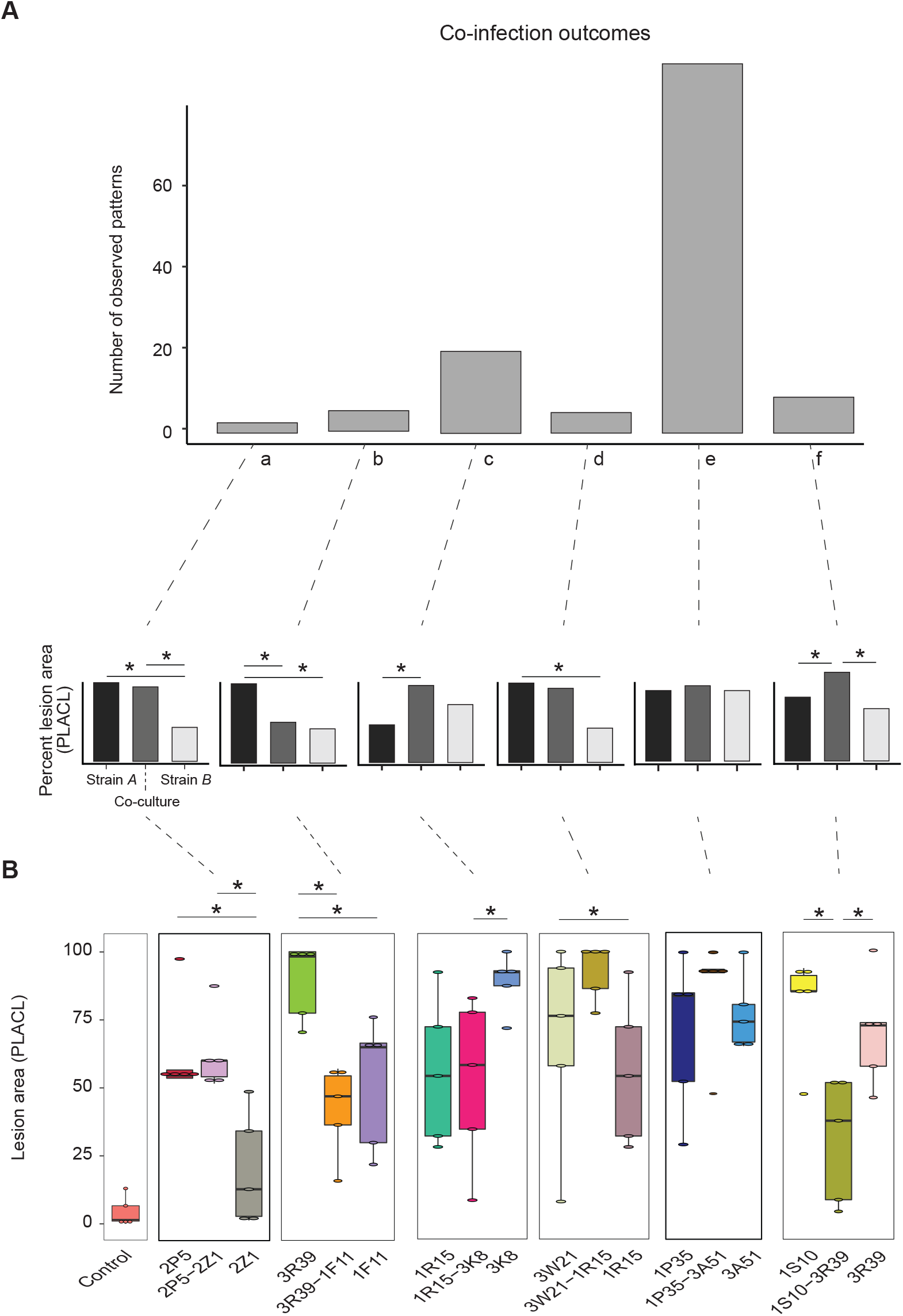
Divergent patterns of co-infection outcomes on wheat leaves. A) Frequency of different pairwise interactions outcomes grouped into six different patterns (a-f). The schematics represent each main pattern. B) Examples of pairwise interactions of wheat single and co-infection. The control represents mock inoculations (significance was assessed using Tukey-Kramer HSD, p < 0.05).

### Contrasting co-culture effects in vitro and during host infections

Given the differences in the environments in which we assessed co-culture and co-infection outcomes, we evaluated the similarity in outcomes of identical strain pairings for a total of 91 matched strain pairings. Single strains showed a moderate but significant correlation between maximum growth rate in culture and the lesion development on wheat leaves (*r* = 0.55, *p* = 0.04; Fig. 5A). Next, we evaluated whether the differences observed between individual strains and the strain mixture were consistent between culture medium and the wheat infection. We found that among all evaluated pairs 30 were found to produce matching interaction patterns and 61 were not matching between the environments (Fig. 5B; for examples see Supplementary Figures S3-4). As an example, the strain 3K8 was among the fastest growing strains in single culture, however 3K8 was not always producing high amounts of lesions on leaves. The strain mixture of 2P5 with 2Z1, the *in vitro* assay showed no growth difference, while infection assay showed mostly more lesion development by 2Z1 compared to 2P5 (Fig. 5C). The interaction between strains 1O40 and 1G30 was similar across conditions with 1O40 being a fast grower *in vitro* and producing high degrees of lesions in the single strain infection assay. The mixture of 1O40 with 1G30 led to both weak growth in culture and less lesions on leaves (Fig. 5C). Finally, we found a majority of mixture assays with divergent outcomes between *in vitro* co-culture and plant co-infection (Fig. 5C). The mixture 1O40-1F11 produced higher growth than any individual strain, however no difference in lesion development was observed among strains and co-infection. In contrast, the mixture 2P5-1G30 showed no significant difference among any single or mixed culture. The same mixture produced more lesions on leaves than any single strain.

**Figure 5:**
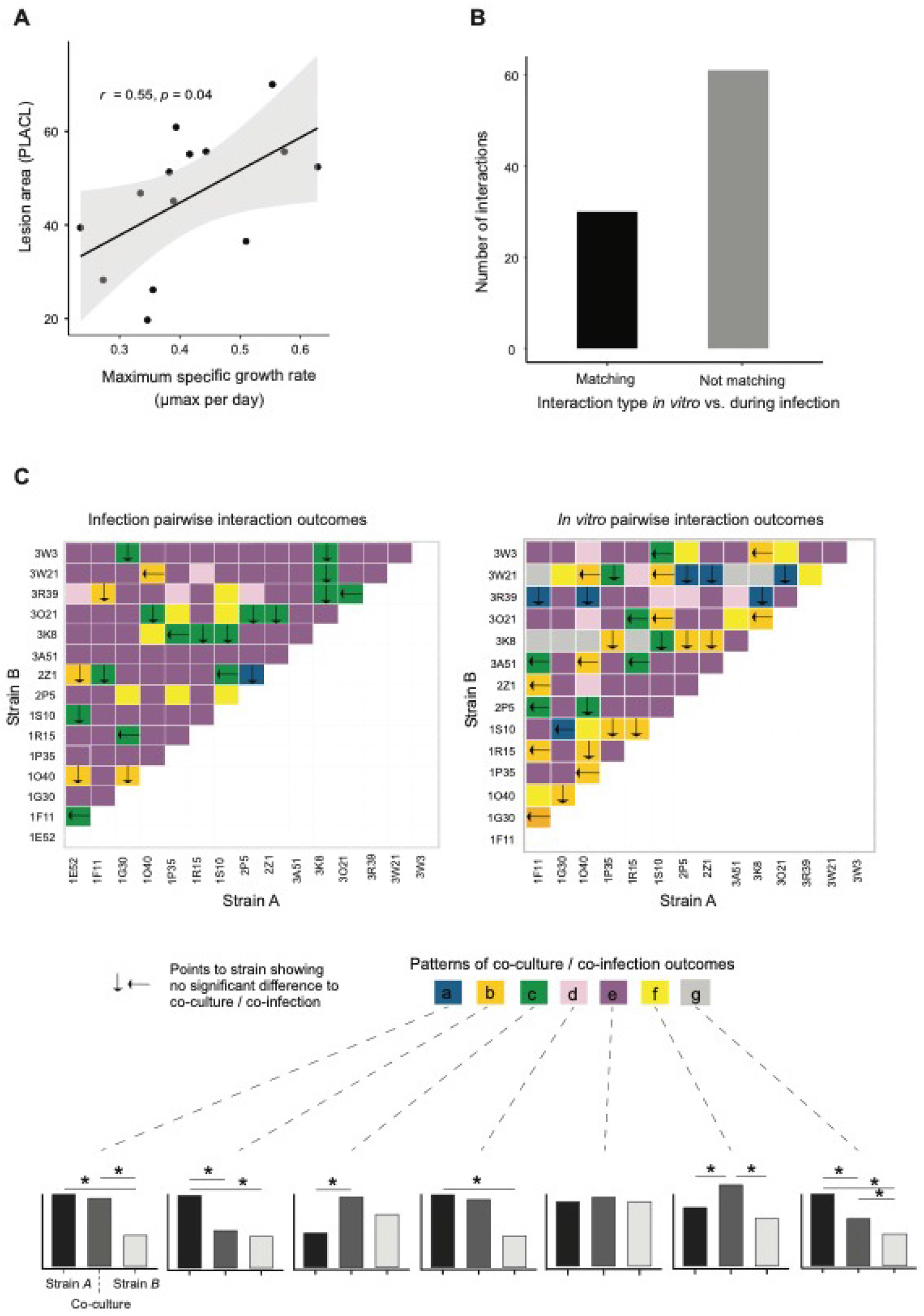
Comparison of in vitro vs. in planta pairwise interaction outcomes. A) Correlation between growth rates of individual strains in culture and lesion development on the plant host. B) Number of matching and mis-matching pairwise interaction outcomes outside and on the plant host. C) Patterns of culture and plant co-infection interaction outcomes for each pair of strains. The schematics at the bottom represent the seven main interaction outcomes (pattern a-g).

## Discussion

Intraspecific co-infections are frequent in nature with plant hosts being often colonized by diverse strain sets (5, 20, 33). Yet, consequences of co-infections are often poorly understood. Our study shows that conspecific genotype pairings can produce divergent outcomes whether the interaction is assessed in culture conditions or as an infection phenotype. Moreover, highly virulent strains are not necessarily the most competitive outside of the host creating the potential for complex selection regimes over the life-cycle of the pathogen.

Competition can be defined as the interaction occurring when organisms accessing a limited resource and having an impact on fitness (34). We found that during single culture experiments outside of the host, individual strains showed heritable differences in their ability to grow on nutrient rich medium consistent with previous studies on *Z. tritici* populations (35, 36). Controlling for the amount of inoculum, co-cultured strains varied widely in their joint growth rates. A large majority of co-cultures produced growth patterns consistent with neutral interactions (*i.e*. no competition). These cases included outcomes where neither the single nor the co-culture showed significant growth differences. Additional cases consistent with this scenario were patterns where the co-culture was showing an intermediate growth with either significant or non-significant differences to each monoculture. For strain pairs showing significant differences in single culture growth, the co-culture was often growing similarly to the weaker strain. However, our experiments lacked the statistical power to clearly distinguish growth differences for most pairings. Despite the large number of tested pairs, we observed no co-culture with significantly reduced growth compared to both monocultures. Hence, strongly suppressive effects among strains appear to be rare under the tested conditions. Interestingly, a small minority of pairings (~5%) showed enhanced growth in the co-culture compared to each monoculture. It is unknown whether such growth acceleration may be triggered by sensing the presence of competitors. An important complement to test hypotheses on mechanisms involved in the competition will be to assess frequencies of individual strains in co-cultures by means *e.g*. of a strain-specific discriminant qPCR or fluorescent labelling techniques. A recent study of four fluorescence-tagged *Z. tritici* strains showed a consistent pattern of fast-growing strains also growing well in co-culture (25), however expanding both labelling and qPCR approaches to much larger pair collections as in this study will be challenging. How fungi recognize self from non-self at different morphological stages and under different environmental conditions remains poorly understood though. Well investigated recognition systems include vegetative incompatibility reactions at the hyphal stage upon non-self contact in *Neurospora crassa* and other filamentous fungi (37, 38). Non-self recognition may be important if strains were to modulate growth investment depending on conspecific competitors.

Strains of *Z. tritici* produced a broad range of lesion symptoms on wheat leaves during infection. We focused on lesion development after a successful infection by the pathogen. The observed range of lesion area per leaf for single strain infections shows that all strains are capable of causing defense responses on the plant host. We observed no meaningful amount of asexual spore structures (*i.e*. pycnidia) in contrast to previous greenhouse experiments with the same pathogen (39). The lack of pycnidia production is most likely related to insufficient inoculation time, host resistance or environmental conditions not conducive for full infections. Nevertheless, the extent of lesion development on leaves is correlated with the success of the pathogen to colonize the leaf surface and trigger defenses. Under field conditions lesion and pycnidia development are correlated (40). Most strain pairs used for co-infections showed no significant differences in lesion development either as a single infection or co-infection. A considerable minority of strain pairs produced intermediate lesion development during co-infection compared to the two strains in single infections. This suggests that under most circumstances co-infections produce only weak deviations in infection outcomes compared to single strain expectations. However, we also found a small set of co-infection pairings where lesion development was accelerated showing that rare co-infections can increase the damage to the plant even at identical inoculum density.

Strain pairs producing faster growth compared to single strains in culture only rarely matched the co-infection effects on lesion development. This mismatch is supported more broadly by having approximately two thirds of all tested co-infection pairs show divergent *in vitro* and plant infection outcomes. Hence, competitive outcomes *in vitro* are only weak predictors for strain interactions on the plant. Such a mismatch between *in vitro* and infection outcomes was also previously observed in an investigation of four *Z. tritici* strains (20). Interestingly, pycnidia spore production (a proxy for fitness) was generally reduced during co-infection compared to single strain infections (20). This points to a potential cost of competition at the level of reproduction. However, larger numbers of strain pairings need to be analyzed under conditions allowing for pycnidia observations to drawer stronger conclusions. Lesion development may only be weakly associated with reproductive output as lesions are largely an expression of host immune responses and damage through reduced photosynthesis. It remains to be investigated how increased lesion development is triggered by the presence of more than one genotype. During the infection process, *Z. tritici* produces hyphae that enter the intercellular space of wheat cells to produce pycnidia in the substomatal space (41, 42). Fluorescent microscopy analyses of co-infections showed that both hyphae and pycnidia are densely packed without apparent exclusion zones among genotypes (25). Increased damage to the host during co-infection could be due to an accelerated hyphal growth of individual strains. Alternatively, different strains may produce different variants of effectors and metabolites during infection (43, 44), which could in combination cause greater damage.

The high frequency at which pathogen strains may be exposed to conspecific strains during infection cycles could create sufficient selection pressure to favor competitive abilities. For instance, selection could favor genotypes retaining growth or infection proficiency despite the presence of competing strains. We identified a small set of strains (*n*=4) retaining growth in each of their pairings with other strains, as well as two strains producing similar amounts of lesions even when exposed to other strains. However, genotyping of co-cultures and co-infections would be necessary to assess how well individual strains reproduce when exposed to other strains. Investigating strategies of pathogens coping frequently with high density environments and exposure to conspecific strains should be an important goal of future co-infection studies. Because co-infections are common in nature (11, 45, 46), understanding the evolution of virulence under co-infection is important for disease management and prevention. Mismatches of competitive outcomes between on and outside of the hosts make predicting co-infection damage more challenging. However, systematic screens of pairwise interactions combined with genetic analyses will help build more informed models of pathogen evolution on crops and other plants.

## Material and methods

### Pathogen strain selection

We used 14 and 15 genetically distinct strains of *Z. tritici* for *in planta* and culture condition experiments, respectively (See Supplementary Table S1). All strains were collected from the Field Phenotyping Platform (FIP) site of the ETH Zürich, Switzerland (Eschikon, coordinates 47.449°N, 8.682°E) in 2016 from 8 winter wheat cultivars (47). After sampling, spores of each strain were stored in either 50% glycerol or anhydrous silica gel at - 80 °C (39). Illumina sequencing datasets are available for all used strains (39).

### In vitro co-cultivation of conspecific strains

*Z. tritici* strains were revived in solid culture media Yeast-Malt-Agar (YMA) added with 50 μg/mL of kanamycin and incubated for 8-10 days in 18°C. A single colony of each strain was selected and inoculated in a 50 ml conical flask containing 20 mL liquid yeast sucrose broth (YSB) medium supplemented with 50 μg/mL of kanamycin. The inoculated flasks were incubated in the dark at 18° C and 140-180 rpm on an orbitary shaker for 8 days. After 8 days of incubation, the cultures were passed through four layers of sterile meshed cheesecloth (Oekostar Textile AG, Switzerland) to eliminate hyphal biomass and retain only spores. Spore suspensions were centrifuged for 15 minutes at 3700 rpm (Vaudaux-Eppendorf AG, Switzerland). After discarding the supernatant, the pellet was washed with sterile water to remove media traces. The process was repeated one more time. Retained spores were grown in 50 mL conical flask, each containing 20 mL of Vogel’s minimal medium supplemented with 50 μg/mL of kanamycin. The flasks were inoculated with spores to reach a spore density of approximately 10^7^ spores per mL. The cultures were then incubated in the dark at 18° C and 140-180 rpm. For growth rate monitoring experiments, spores were filtered, diluted in fresh minimal media, and the cell density adjusted to 10^5^ spores/mL by counting the cells in a KOVA cell chamber system (Kova International, USA). Strains were then cultured alone or in pairs in 96-well plate in either 100 μL or 200 μL, respectively. The plates were sealed and incubated at 18 °C at 12 rpm on a shaker incubator for 11 days with optical density measurements (OD600) once daily on a SpectraMax i3x plate reader (Molecular Devices, USA) with a shaking period of 5 s prior to the measurement. The OD was exported to Microsoft Excel for further analysis and generation of the growth curves. At least 5 replicates for each single or paired culture were set up (See Supplementary Table S3).

### Wheat cultivar selection and infection assays

The wheat (*Triticum aestivum L*.) cultivar Combin was used due to its susceptibility to *Z. tritici* (48). Five seeds of the cultivar Combin were placed in pots filled with compost soil and placed in white watering boxes. The pots were set to replicate (*n*=5) each treatment of single and co-infection. Wheat was cultivated during 21 days in a growth chamber with a photoperiod of 16h light / 8h dark, a temperature of 18°C and a relative humidity of at least 60%. Plants were watered when needed and randomized regularly. Strains of *Z. tritici* used in the infection experiment were revived from stock in solid culture media yeast malt agar (YMA) supplemented with 50 μg/mL of kanamycin as described above. Strains were cultured at a temperature of 18° for 8-10 days. Following this time period, a single colony of each strain was selected and used to inoculate 20 mL of liquid culture media containing yeast sucrose broth (YSB) supplemented with 50 μg/mL of kanamycin. The incubation was at 18°C on a shaker for 8-10 days at 160 rpm. Following this step, strains were filtrated using sterile cheese cloth (Oekostar Textile AG, Switzerland) and placed in a cold room at 4°C. Then, 3-5 days before their use in the experiment, the strains were taken from the cold room and 500μL of each strain was used to inoculate 20 mL of liquid YSB and placed in a shaker at 140-160 rpm and 18°C. The strain cultures were filtered using sterile cheese cloth (Oekostar Textile AG, Switzerland) and centrifuged for 15 min at 3700 rpm (Vaudaux-Eppendorf AG, Switzerland). After discarding the supernatant, the pellet was resuspended in 20 mL of Milli-Q water (Merck Group, Switzerland) and centrifuged again for 15 min at 3700 rpm. This step was repeated once more. The spore concentration of each strain was adjusted to either 10^5^ spores per mL or half the concentration for co-infections using a hemocytometer (Kova International, USA). The strains were diluted in Milli-Q water (Merck Group, Switzerland) to reach the desired concentrations. Finally, 0.1% of Tween 20 (Sigma-Aldrich, Switzerland) was added to facilitate wheat leaf spore adhesion.

Treatments consisted of plants sprayed with water only (control treatment), plants infected with one strain at 10^5^ spores per mL (single strain control), or plants co-infected with two strains with a concentration of 5 * 10^4^ spores per mL per strain (co-infection treatment). After, twenty-one days of seed sowing, each pot was sprayed separately protected by cardboard to prevent splashing. Sprayers containing either spore suspensions or sterile Milli-Q water (Merck Group, Switzerland) were used. Each pot was placed in white container boxes and sealed with plastic bags and adhesive tap after being watered. The sealing helps maintaining 100% humidity in a room at 18°C for 48 hours. After this period, each plastic bag was opened and left in the climate-controlled room for 21 days. The plants were watered when needed and pots were regularly randomized within each white box. White boxes were also regularly randomized.

### Plant infection assessments

After twenty-one days of infection, the first two leaves of each plant were cut and pasted on paper sheets containing a QR code for tracking. Then, each sheet was scanned at 1200 dpi using a flatbed scanner (EPSON perfection V55O). Following this step, an automated image analysis was performed using ImageJ (Rasband, W.S., ImageJ, U. S. National Institutes of Health, Bethesda, Maryland, USA, https://imagej.nih.gov/ij/, 1997-2018). We used a macro developed to assess *Z. tritici* disease symptoms (47) to determine the percentage of leaf area covered by lesions (PLACL), which is a proxy for leaf damage and virulence (49). Automatic assessments were visually checked for consistency. PLACL values were adjusted if color thresholds were not appropriately distinguishing yellowish (*i.e*., lesion areas) and green (*i.e*., healthy) areas on some leaves. Overall, very little pycnidia formation was observed during the infection period and pycnidia counts were, therefore, not considered further.

### Data analyses

All data analyses were carried out using R v4.0.4 software (50). Optical densities were corrected for the blank control (Vogel’s minimal media) by subtracting the mean optical density for each strain. Fungal growth curves were estimated by fitting a log-logistic model to the data using the {Growthcurver} R package (51). Furthermore, {Growthcurver} was used to summarize the following growth characteristics: generation time (td) and the maximal specific growth rate (*μmax*). Visualizations were made using the {ggplot2} R package (52).

Analysis of single versus co-culture data was performed using one-way ANOVA and Tukey-Kramer tests for post-hoc analysis in R. To satisfy normality assumptions, PLACL data was square-root arcsine transformed. Data handling was performed with the R packages {dplyr} (53), {reshape2} (54) and {tidyr} (55). We used a linear model to test the effect of the different treatments on PLACL, followed by a one-way ANOVA with the function “Anova” from the {car} package (56). Assumptions of linear models were visually assessed with the function “autoplot” from the package {multcomp} (57). Post-hoc analysis was done using the “emmeans” function from the package {emmeans} (58) with the Tukey-Kramer method for multiple testing correction.

## Data availability

All generated data is provided in the supplementary material.

## Acknowledgements

We thank Nikhil Kumar Singh, Luzia Stalder and Guido Puccetti for advice on plant infection protocols and in vitro growth experiments. We also thank Laetitia Holzer for her assistance with lab work and helpful discussions.

