## Supplementary Figures for "Divergent outcomes of direct conspecific pathogen strain interaction and plant co-infection suggest consequences for disease dynamics"

**Supplementary Material**

**Supplementary Figures**

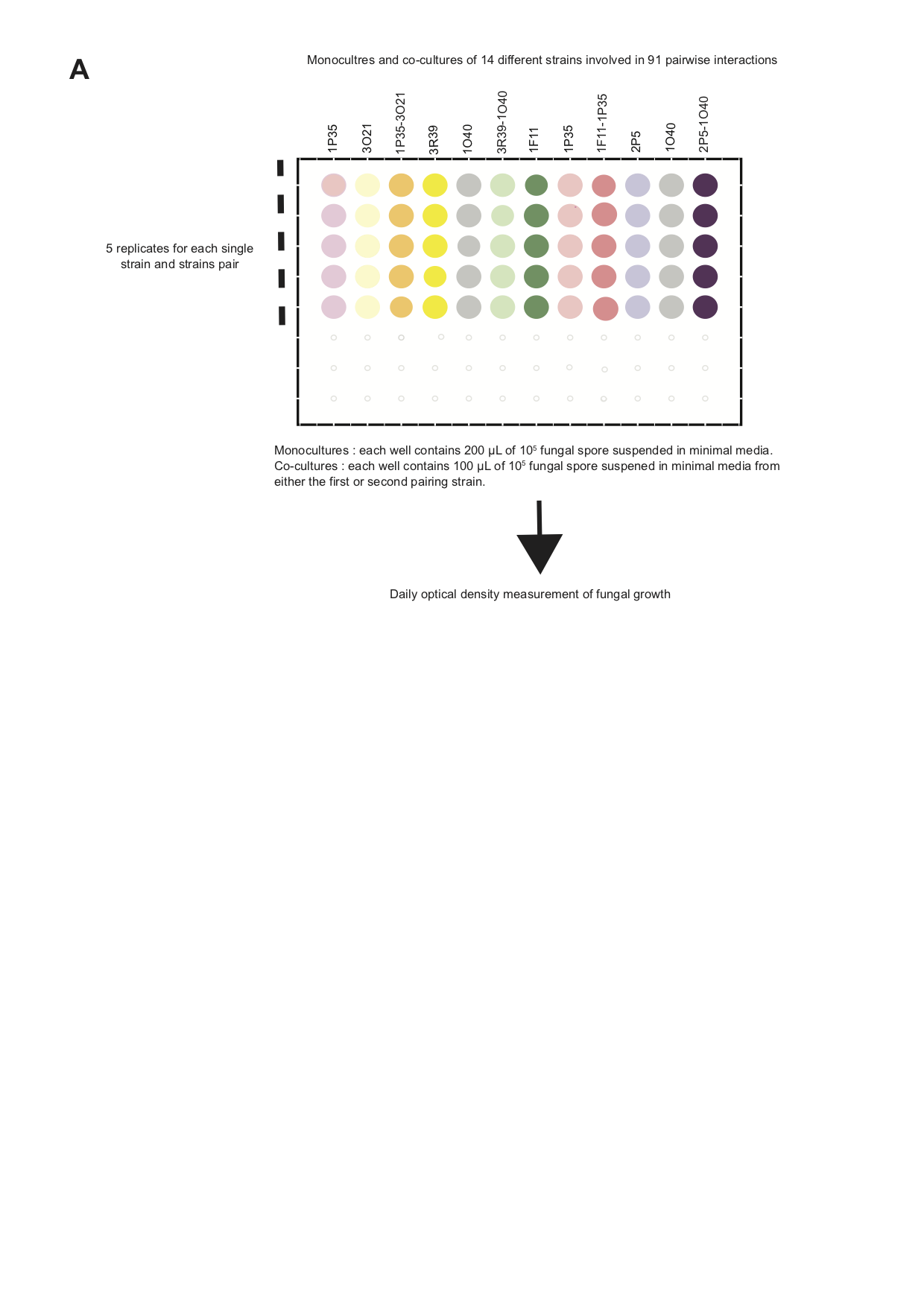

**Supplementary Figure S1: Schematic of the *in vitro* culture growth assessment in microtiter plates**. A) Schematic overview of in vitro fungal strains growth assessment during single and mixed culture using optical density (OD) measurements. For each monoculture a total volume of 200 μL of fungal spore concentration of 10^5^ suspended in minimal media was cultured in each microplate well, whereas for mixed culture, each microplate well was containing 100 µL of fungal spores each from two different strains. At least 5 replicates for each single or paired culture were set up.

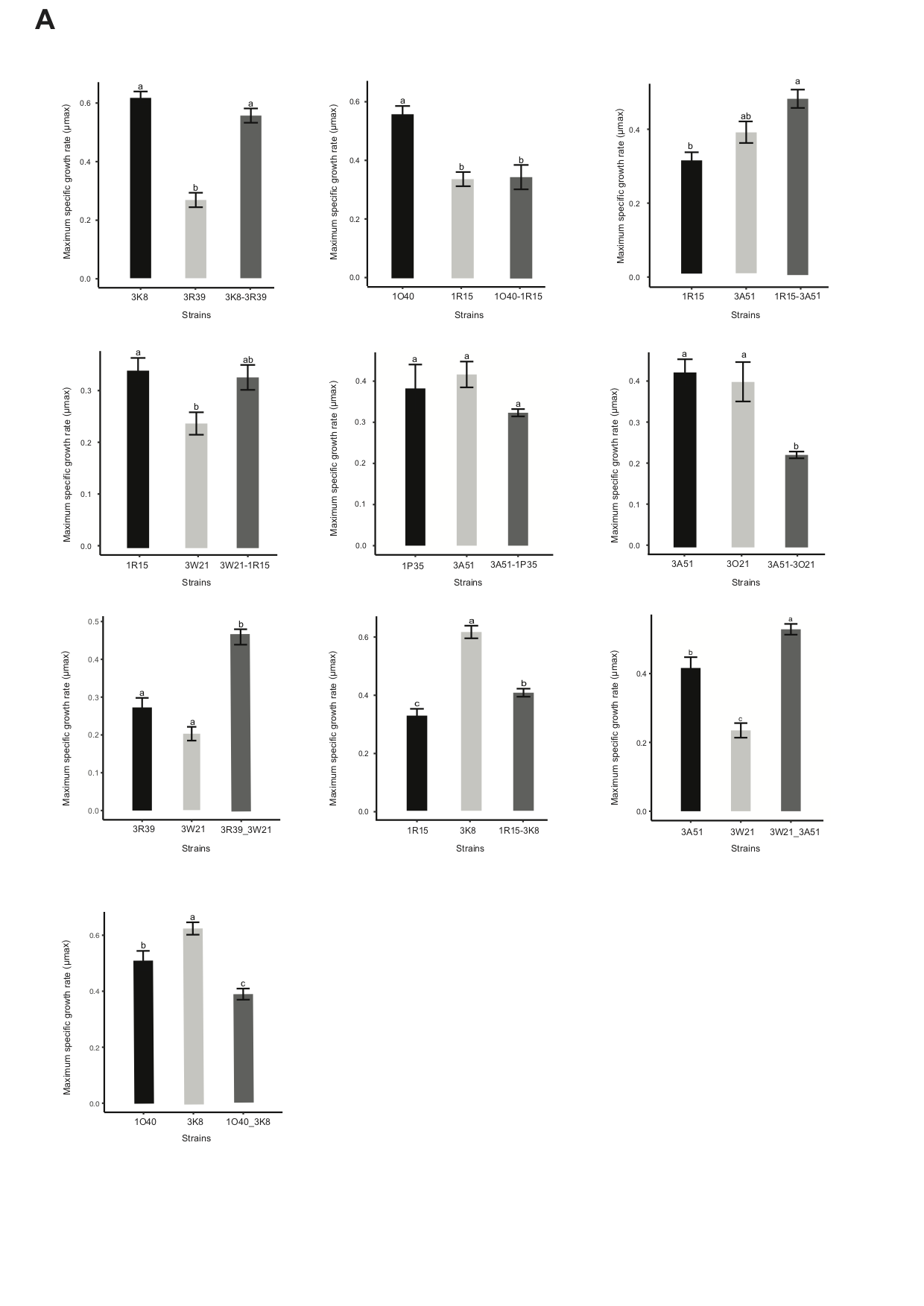

**Supplementary Figure S2:** Examples of pairwise strain interaction outcomes for *in vitro* cultures.

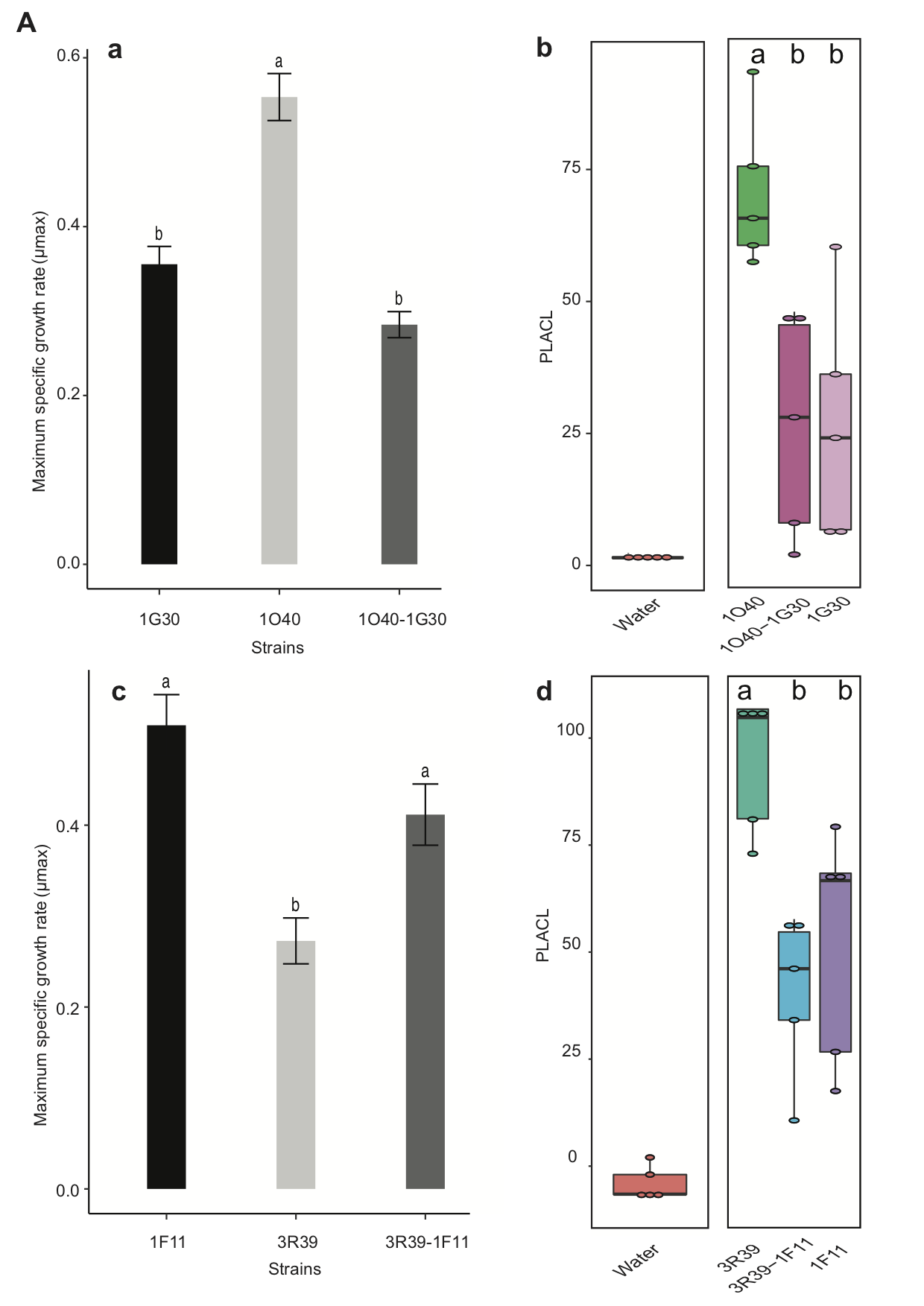

**Supplementary Figure S3:** Examples of pairwise strain interaction outcomes in culture and on the plant host.

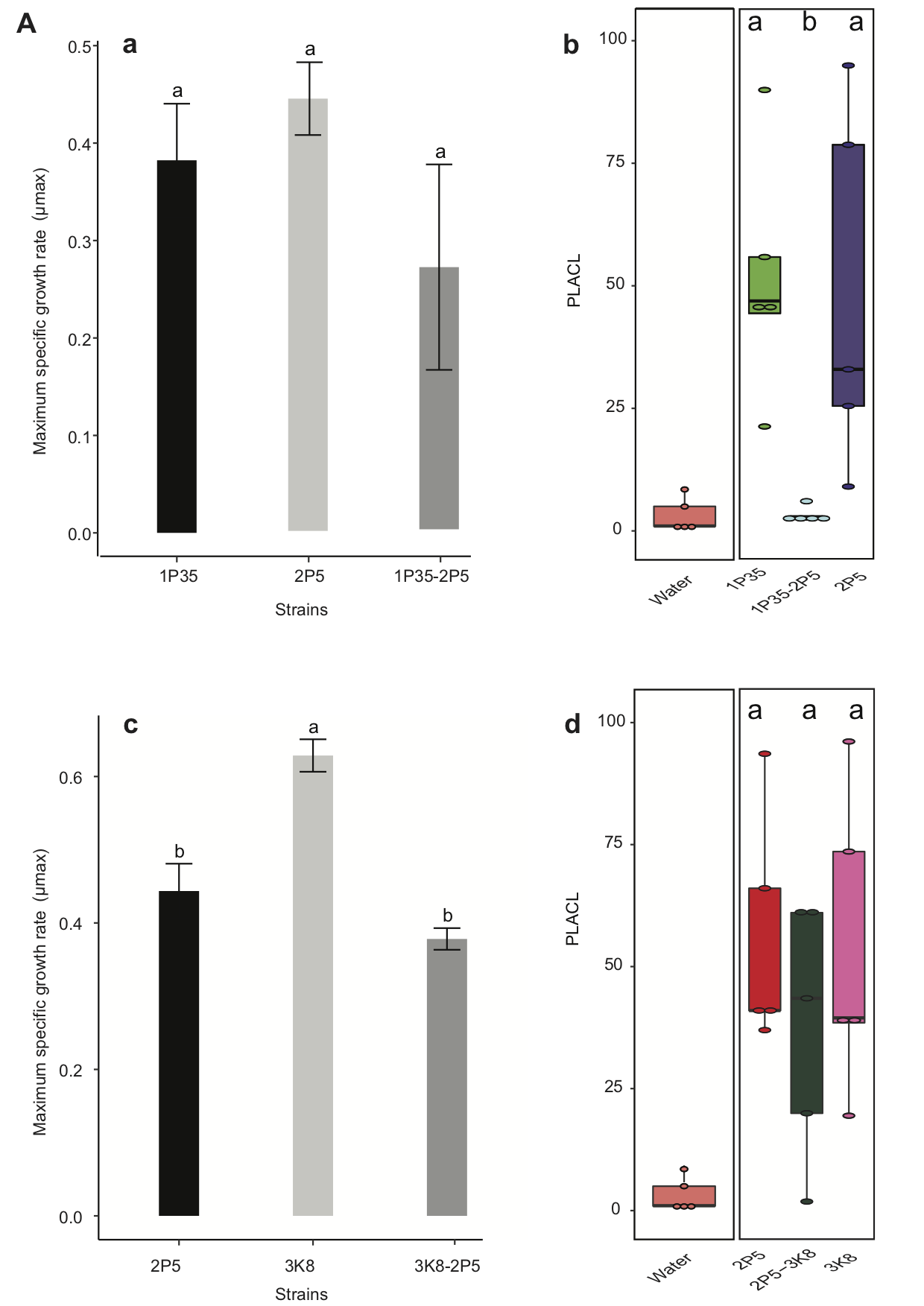

**Supplementary Figure S4:** Examples of pairwise strain interaction outcomes in culture and on the plant host.

**Supplementary Table legends**

(See Excel file)

**Supplementary Table S1:** Strains used either *in vitro* or for plant pairwise interaction experiments.

**Supplementary Table S2:** Identification of strain pairs tested either *in vitro* or *in planta*.

**Supplementary Table S3:** Optical density (OD) measurements and estimates of growth metrics.

**Supplementary Table S4:** Percent leaf area covered by lesions (PLACL) measurements for infection experiments.
